# Integrated Systems Analysis Deciphers Transcriptome and Glycoproteome Links in Alzheimer’s Disease

**DOI:** 10.1101/2023.12.25.573290

**Authors:** Yusuke Matsui, Akira Togayachi, Kazuma Sakamoto, Kiyohiko Angata, Kenji Kadomatsu, Shoko Nishihara

## Abstract

Glycosylation is increasingly recognized as a potential therapeutic target in Alzheimer’s disease. In recent years, evidence of Alzheimer’s disease-specific glycoproteins has been established. However, the mechanisms underlying their dysregulation, including tissue- and cell-type specificity, are not fully understood. We aimed to explore the upstream regulators of aberrant glycosylation by integrating multiple data sources using a glycogenomics approach. We identified dysregulation of the glycosyltransferase PLOD3 in oligodendrocytes as an upstream regulator of cerebral vessels and found that it is involved in COL4A5 synthesis, which is strongly correlated with amyloid fiber formation. Furthermore, COL4A5 has been suggested to interact with astrocytes via extracellular matrix receptors as a ligand. This study suggests directions for new therapeutic strategies for Alzheimer’s disease targeting glycosyltransferases.

**Graphical Abstract:** 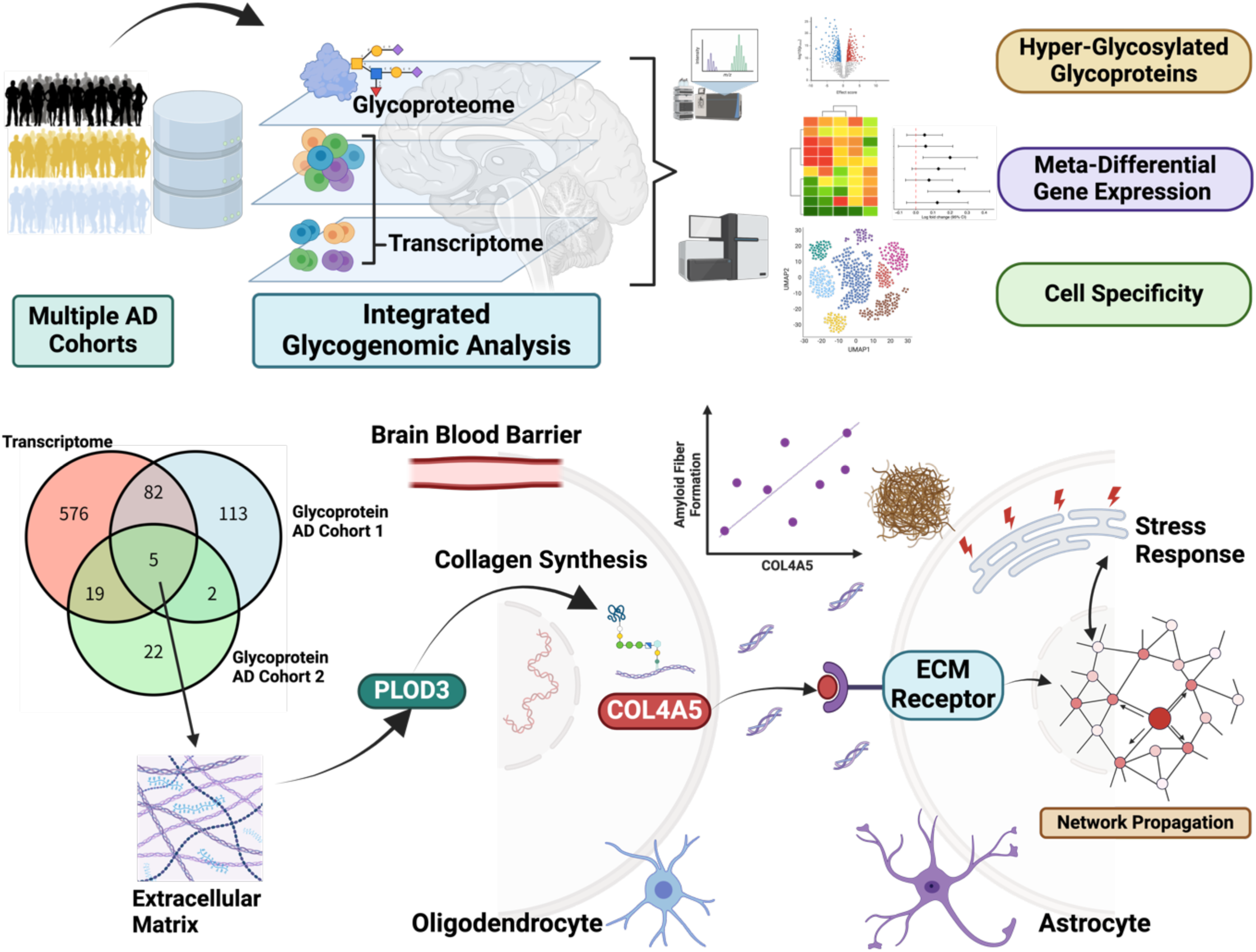

## Introduction

Alzheimer’s disease (AD) is an age-related neurodegenerative disease^1,2^. The primary causes are neurogenic cell loss, accumulation of misfolded proteins, oxidative stress, and inflammatory responses^3^. Genomic, transcriptomic, and epigenetic mechanisms have been intensively examined^4^. However, our knowledge of the post-translational modifications that regulate cellular functions and interactions between cells is still lacking^5^. In particular, glycosylation is the most diverse and abundant post-translational modification among protein modifications^6^.

Protein glycosylation is a complex multistep process involving approximately 200 different glycosyltransferases^6–8^. There are 16 major glycosylation pathways, including lipid glycosylation, N-glycosylation, O-glycosylation, C-mannosylation, lipid glycosylation, and glycosylphosphatidylinositol (GPI)-anchored synthesis. Recently, glycomics / glycoproteomics analysis of the human AD postmortem brain^9,10^, serum,^11–17^ and cerebrospinal fluid^18–20^ for N-glycosylation, the most abundant glycosylation pathway, has revealed dysregulated glycoproteins. In addition, the biological functions of abnormal glycans in AD pathology have been reported in some cases; for example, it is known that inhibition of BACE1 glycosylation reduces the cleavage of the amyloid β precursor protein (APP) ^21–23^. However, most biological functions of glycosylation in the pathogenesis of AD are poorly understood.

Glycan structures are not independent of the DNA template, and glycosylation depends on a combination of approximately 200 glycosyltransferases and 500 related proteins^6–8^. Thus, their dysregulation may act as an upstream regulatory factor that triggers abnormal glycosylation processes^24^. In addition, it is difficult to elucidate biological glycosylation mechanisms at the single-cell resolution using glycomics / glycoproteomics alone because current technology is limited to probing with glycosylation-specific antibodies and glycan-binding proteins, such as lectin ^6^. Therefore, a glycogenomics approach that integrates genomics, or functional genomics, and glycoproteomics is critical for a comprehensive understanding of biological glycosylation pathways^24,25^.

Here, we present the factors upstream of aberrant glycosylation in AD. We performed an integrated analysis of bulk and single-cell/nuclear transcriptomic and glycoproteomics data from human AD brain tissues. In particular, we showed that the extracellular matrix (ECM) is a common signature in the glycoproteome and transcriptome, and that ECM gene expression signatures are enriched for cerebral vascular-related pathways. We identified Procollagen-Lysine,2-Oxoglutarate 5-Dioxygenase 3 (PLOD3) as an upstream glycosyltransferase common to ECM pathway. Through an integrated analysis of multiple single-cell expression data, we showed that PLOD3 is involved in the regulation of collagen type IV alpha 5 chain (COL4A5), which is strongly correlated with amyloid fiber formation. Cell–cell interaction and signaling pathway analyses suggested that PLOD3-COL4A5 cascade is involved in the stress response via the ECM receptor in astrocytes.

## Results

### Hyperglycosylated proteins are primarily enriched in the ECM

To examine the association between the molecular pathogenesis of AD and glycosylation, we accessed glycoproteomic data consisting of 2 cohorts of postmortem brain tissue from AD patients ^9,10^ (**Figure 1A**). The first dataset consisted of dorsolateral prefrontal cortex tissue from 8 neuropathologically confirmed AD cases and 8 age-matched controls. The second dataset comprised a subset of the ROSMAP cohort^14^. Glycoproteomic analysis was performed on the postmortem brains of 10 patients with asymptomatic AD, 10 patients with symptomatic AD, and 10 healthy brains in which none of the above were present.

**Figure 1.**
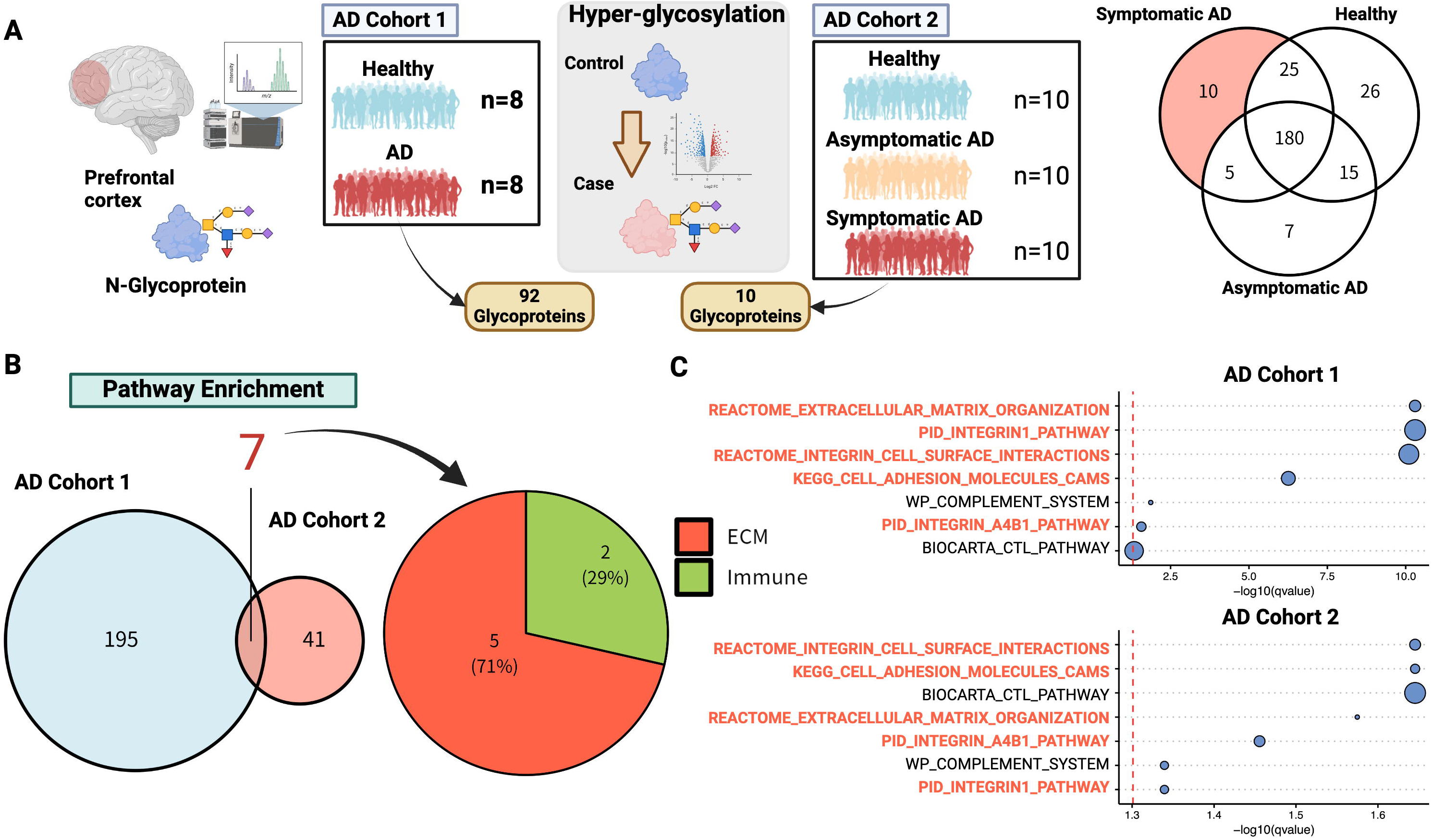
Hyperglycosylated proteins are primarily enriched in the ECM. A Analysis of glycoprotein data from 2 AD cohorts, using glycoproteins from prefrontal tissues of 2 independent AD cohorts. The first cohort (AD cohort 1) consisted of 8 samples each from healthy subjects and those with AD, and the second cohort (AD cohort 2) consisted of 10 samples each from healthy subjects, asymptomatic AD, and symptomatic AD. In each cohort, 92 and 10 AD-specific glycoproteins were identified, respectively. B Pathway enrichment of AD-related glycoproteins. Over-representation analysis of AD-specific glycoproteins was performed. C Significantly enriched pathways that were common in both cohorts are shown. The horizontal axis is the p-value representing the enrichment, which is the logarithm of the nominal p-value multiplied by a negative value. Figures were created with BioRender.com.

In each cohort, 92 and 10 AD-specific hyperglycosylated proteins (**Supplementary Table S1**) were identified, respectively, and pathway enrichment analysis was performed (**Figure 1A**). Among the pathways significantly enriched in the 2 cohorts, we identified the ECM pathway as the most common pathway among 7 pathways (**Figure 1B, C**). The relationship between AD and ECM has recently been recognized as a new molecular pathogenesis, along with other major pathological hypotheses^26,27^. ECM components contain glycoproteins, including glycosylated proteoglycans and collagen, as major elements^27^, and many glycosylations play important roles in ECM formation and maintenance.

### Meta-analysis of the transcriptome reveals that glycogenes are enriched in the ECM

We explored the upstream factors that regulate ECM hyperglycosylation in AD. We accessed the AD Knowledge Portal (https://adknowledgeportal.synapse.org), which contains postmortem brain transcriptome data from multiple cohorts of patients with AD, and compiled gene expression data. The glycogene set consisting of 214 glycosyltransferases was defined using the gene list in the Glycogene Database (GGDB: https://acgg.asia/ggdb2/)^28^ and literature^6,29,30^ (**Figure 2A, Supplementary Table 2**). This gene set was also categorized according to its glycosylation pathway and synthesis steps (**Figure 2A**).

**Figure 2.**
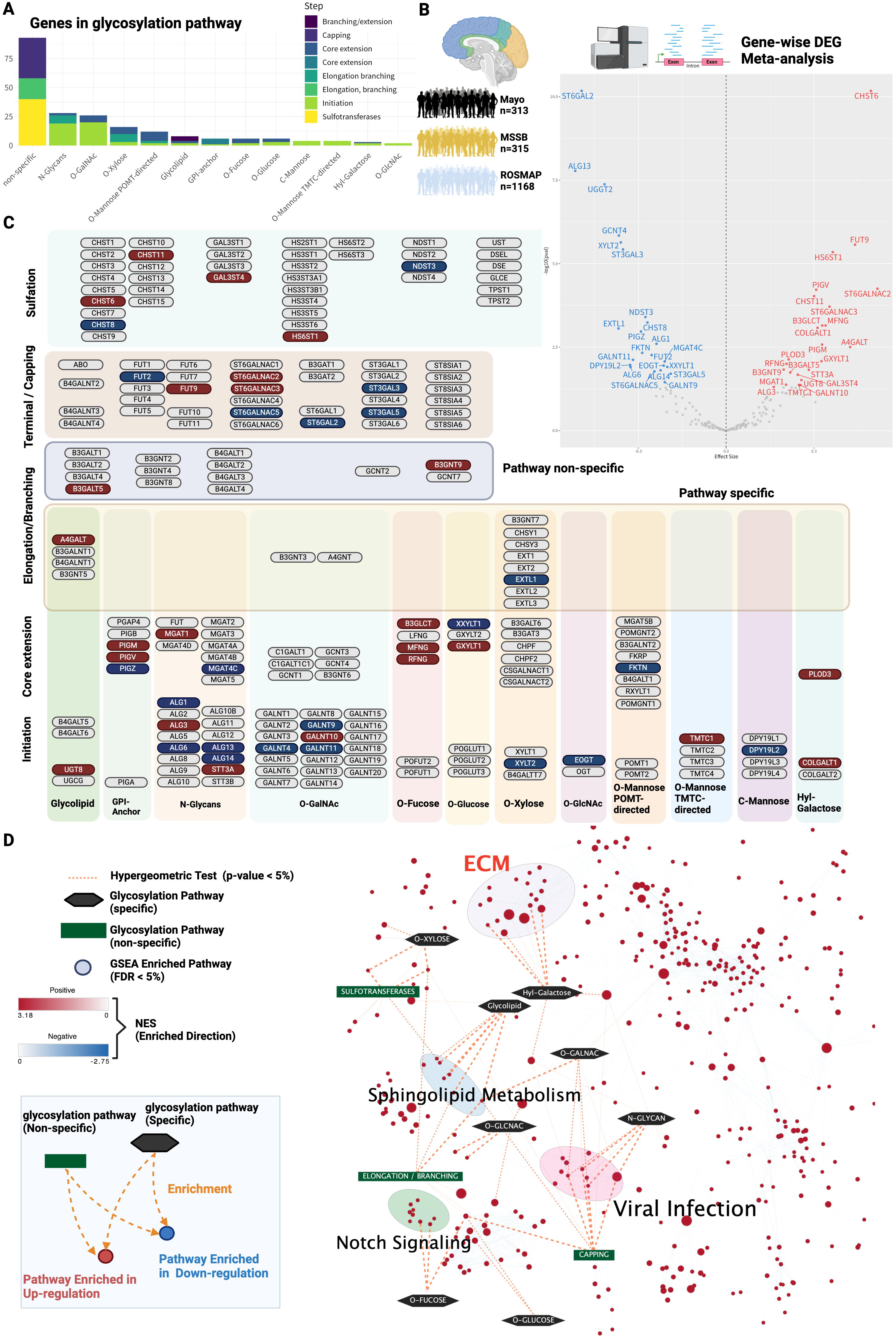
Meta-analysis of the global transcriptome reveals that glycogenes are enriched in the ECM. A Number of glycogenes constituting the glycosylation pathway used for transcriptome analysis. B Meta-analysis of differential gene expression in multiple AD cohorts. Transcriptome data from 3 AD cohorts: the Mayo cohort (n = 313), the MSSM cohort (n = 315), and the ROSMAP cohort (n = 1168). A meta-analysis of DEGs based on gene-level expression levels (FDR < 5%) was performed; 46 glycogenes were identified as DEGs. In the volcano plot, the horizontal axis represents the effect size summarizing the difference in expression between the non-AD and AD groups across cohorts, and the vertical axis represents the log of the p-value from the meta-analysis (bottom is 10) multiplied by a negative value. C Mapped glycosyltransferase DEGs. In total, 46 glycogene DEGs were mapped. Genes overexpressed in the meta-analysis are shown in red, genes underexpressed are shown in blue, and genes that did not show significant mutations are shown in gray. Genes are classified into 16 major glycosylation pathways, including initiation, core elongation, elongation/branching, capping, and sulfation. Glycosyltransferases with and without pathway specificity are also distinguished. Figure created by BioRender.com.

We derived transcriptional signatures of glycogenes based on a meta-analysis. We identified 46 differentially expressed genes (DEGs) in the glycogenes (**Figure 2B, Table S3**). We mapped glycogenes to glycosylation pathways to determine the pathways enriched for the DEGs (**Figure 2C**). Glycosyltransferases were differentially expressed in all pathways (**Figure 2C**), indicating that the signals triggering aberrant glycosylation had already been observed at the transcriptional level.

Next, we analyzed the biological functions of these glycogene signatures. The 779 globally enriched biological pathways were estimated based on the effect size from the differential expression obtained by meta-analysis using all genes (FDR < 5%) (**Figure 2D, Supplementary Table S2–S4**). Subsequently, a post-hoc enrichment analysis was performed to infer which glycosylation pathways were associated with these enriched biological pathways (**Figure 2D, Supplementary Table S5**). Significant glycosylation pathways were extracted with a hypergeometric test as the final estimation results (**Figure 2D, Table S4–S5; p-value < 5%**). We found that the ECM is a common biological signature of the transcription and glycosylation layers in AD (**Figure 3A**). The ECM cluster was strongly associated with the hydroxyl galactose glycosylation pathway (**Figure 2D**).

**Figure 3.**
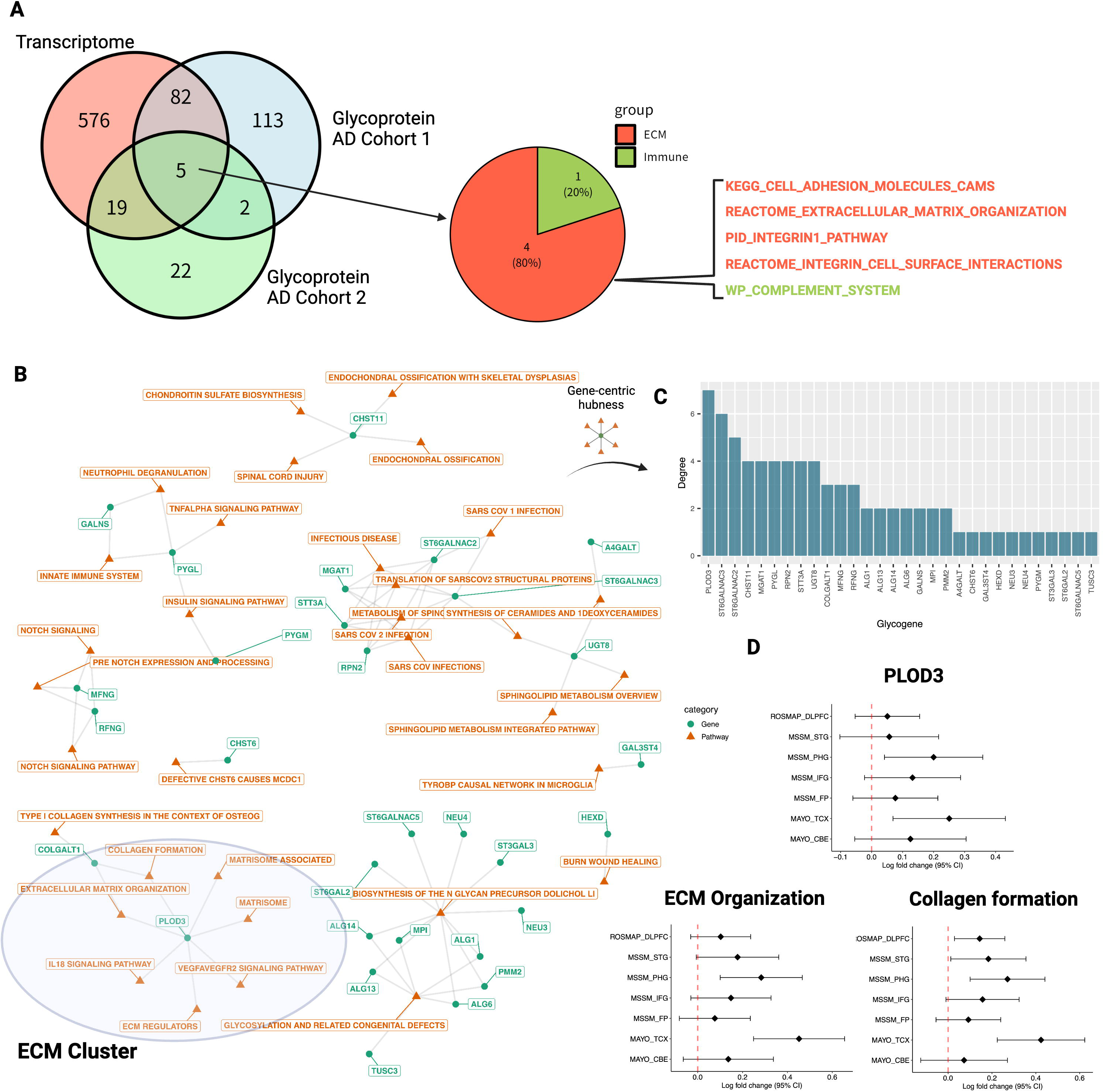
PLOD3 is identified as a hub glycogene for the ECM. A Comparison of AD trascriptome and glycoprotein signatures. Common pathways are shown. B Relationship between glycogenes and globally enriched pathways. Orange nodes represent globally enriched pathways, and green nodes represent glycogen enriched in each pathway. C Functional hub glycogenes in globally enriched pathways. To identify functional hub glycogenes involved in multiple pathways, we constructed a pathway–gene bipartite graph, calculated the degree of each glycogene (number of genes directly connected to the pathway), and ranked the importance of each glycogene. The vertical axis of the bar graph represents the order of each glycogene. D Activity changes in PLOD3, ECM, and collagen formation in AD brains in each transcriptome cohort. Forest plots of Log2 fold changes in PLOD3, ECM organization, and collagen formation activity between non-AD and AD are plotted by cohort and brain region. DLPFC: dorsolateral prefrontal cortex; STG: superior temporal gyrus; PHG: parahippocampal gyrus; IFG: inferior frontal gyrus; FP: frontal pole; TCX: temporal cortex; CBE: cerebellum. Dots indicate estimated mean effect sizes, bar widths are 95% confidence intervals of the estimates, and vertical lines with red dots indicate zero (no change).

### PLOD3 is identified as a functional hub glycogene for ECM

Next, we performed an in-depth analysis of the glycogenes that play a central role in the ECM. Of the 779 globally enriched pathways, we constructed a bipartite graph consisting of glycogene–pathway relationships based on 48 pathways, including the differentially expressed glycogenes (**Figure 3B**). We inferred the glycogene importance based on the number of neighboring pathways, that is, the network degree (**Figure 3C**). As a result, PLOD3 was identified as a hub glycogene with the highest degree (**Figure 3C**).

PLOD3 is an enzyme that mediates essential glycosylation during the early stages of collagen formation^31^. In general, collagen is broadly modified by the hydroxylation of proline and lysine and glycosylation of specific hydroxylysine residues^32^. Hydroxylation of lysine is catalyzed by PLOD3^33,34^; hydroxylysine undergoes further glycosylation, and COLGALT1 transfers galactose, which are critical steps for maintaining collagen integrity^32^.

To further confirm the results at the gene expression level, we examined whether the changes in PLOD3 expression were consistent among the AD cohorts included in the meta-analysis. We found that PLOD3 was consistently upregulated in individual cohort studies (**Figure 3D**), and the expression signatures of ECM organization and collagen formation showed a consistent overexpression trend (**Figure 3D**). Based on this analysis, we hypothesized that hyperglycosylation of the ECM in AD brain tissue is mediated by PLOD3.

### PLOD3 is expressed in oligodendrocytes and co-expressed with COL4A5

We sought to determine the cellular origin of the PLOD3 and collagen genes. First, we accessed the scRNA-seq data of normal brain tissue from the Human Protein Atlas (v22)^35,36^. We found that PLOD3 was co-expressed with COL4A5 in oligodendrocytes (**Figure 4A**). These two genes showed distinct oligodendrocyte-specific expression signatures (**Figure 4B**). We also accessed a human AD cohort of single-nucleus RNA-seq data for the entorhinal cortex (GSE138852)^37^. The entorhinal cortex is one of the brain regions that shows neurodegeneration in the early stages of AD^38–40^. The cohort included both non-cognitive impairment (NCI) and AD brain. Six cell types were identified: microglia, astrocytes, neurons, oligodendrocyte progenitor cells, oligodendrocytes, and endothelial cells (**Figure 4C**). PLOD3 and COL4A5 were highly expressed in oligodendrocytes (**Figure 4C**). These genes were also predominantly expressed in the AD group (**Figure 4C**).

**Figure 4.**
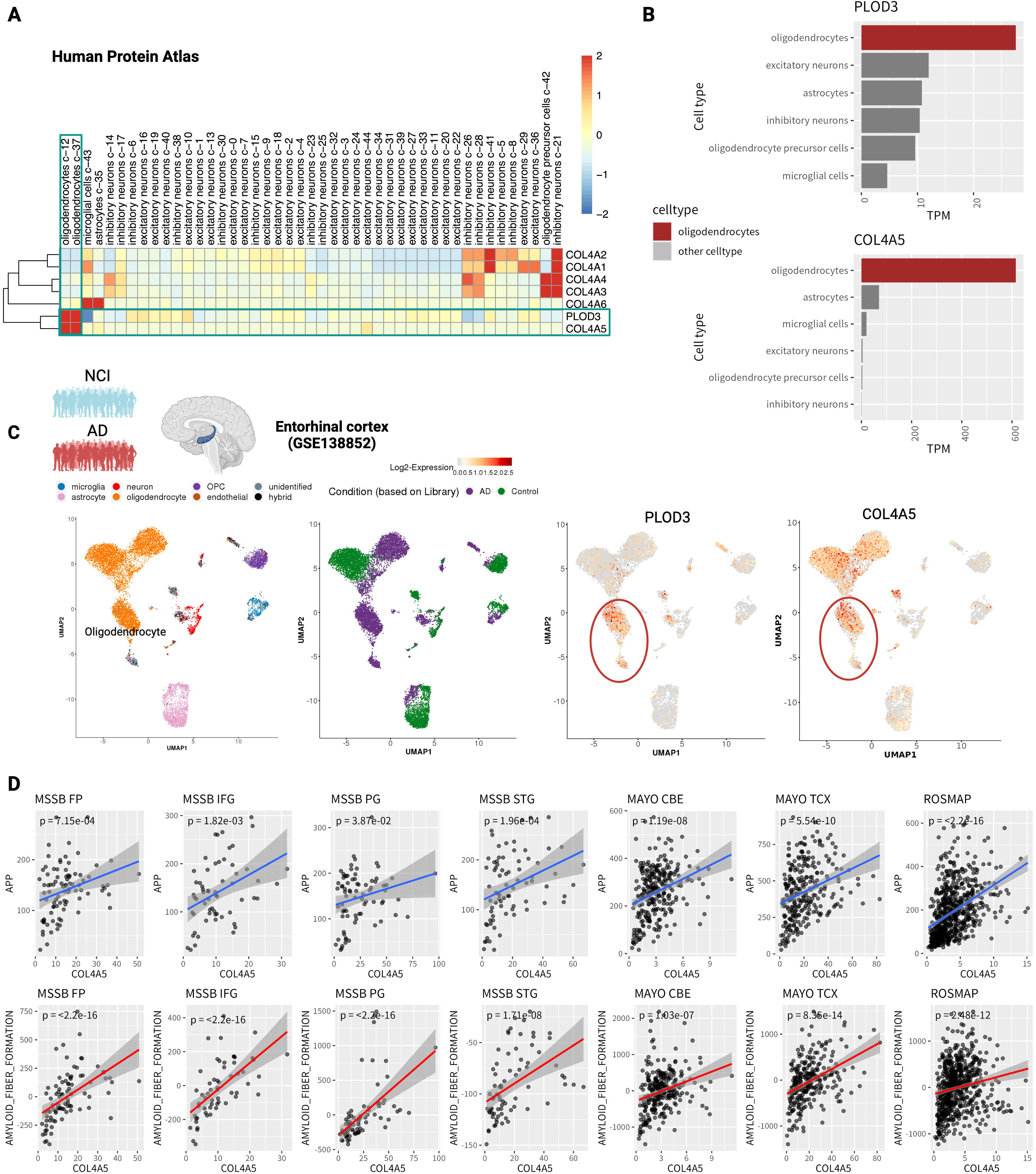
PLOD3 is expressed in oligodendrocytes and is co-expressed with COL4A5. A Cell-type specificity of PLOD3 in healthy brain tissues. Cell clusters obtained from gene expression in healthy brain tissue by Human Protein Atlas (v22) scRNA-seq, and the Transcripts Per Million (TPM) in each cluster. PLOD3 and COL4A5 are highly expressed in oligodendrocytes and belong to the same cluster. B Expression levels of PLOD3 and COL4A5 per cell type. C Cellular specificity of PLOD3 and collagen in the enthorhinal cortex. Scatter plots show the cluster structure of cell populations projected by UMAP to 2D coordinates based on gene expression; the first panel shows cell types, the second non-AD and AD, the third and fourth panels show cell type-specific expression of PLOD3 and COL4A5 in oligodendrocytes, respectively. D Correlation of COL4A5 with the expression of APP (upper panel) and the activity of amyloid fiber formation (lower panel) for each cohort and each region. DLPFC: dorsolateral prefrontal cortex; STG: superior temporal gyrus; PHG: parahippocampal gyrus; IFG: inferior frontal gyrus; FP: frontal pole; TCX: temporal cortex; CBE: cerebellum.

### COL4A5 consistently correlated with amyloid fiber formation in multiple cohort studies

COL4A5 is partially correlated with amyloid plaque accumulation^41^. However, this finding has not been validated in large clinical samples. We tested whether COL4A5 expression was significantly correlated with APP expression. We analyzed the bulk RNA-seq data used in the meta-analysis and examined their relationship with APP gene expression separately for each brain region. The results showed that COL4A5 was strongly correlated with the APP gene in all datasets (**Figure 4D**). Furthermore, we defined the gene signatures of the amyloid plaque formation pathway and analyzed the correlation between their eigengene expression and COL4A5 in the same way. As expected, a strong correlation was confirmed (**Figure 4D**). PLOD3 was evaluated similarly, showing a weaker correlation than COL4A5, but it was significant in several datasets (**Supplementary Figure 1**).

### Cerebrovasculature most strongly associated with ECM dysregulation

We explored whether overexpression of the PLOD3–COL4A5 axis is involved in biological processes in the AD brain. First, we analyzed the biological pathways that best explained ECM activity. We used AES-PCA^42–44^, a principal component analysis (PCA)-based regression model with ECM activity as the outcome variable and all other biological pathway activities as predictors, for each AD cohort used in the meta-analysis (**Figure 5A, Supplementary Table S6**). The estimated p-values were statistically combined using Fisher’s method (**Figure 5A**). Four of the top 10 enriched genes were associated with the vascular system (**Figure 5A**) and were overexpressed in the AD group at the expression level (**Figure 5B**). We hypothesized that the PLOD3–COL4A5 axis is involved in the cerebrovascular microenvironment.

**Figure 5.**
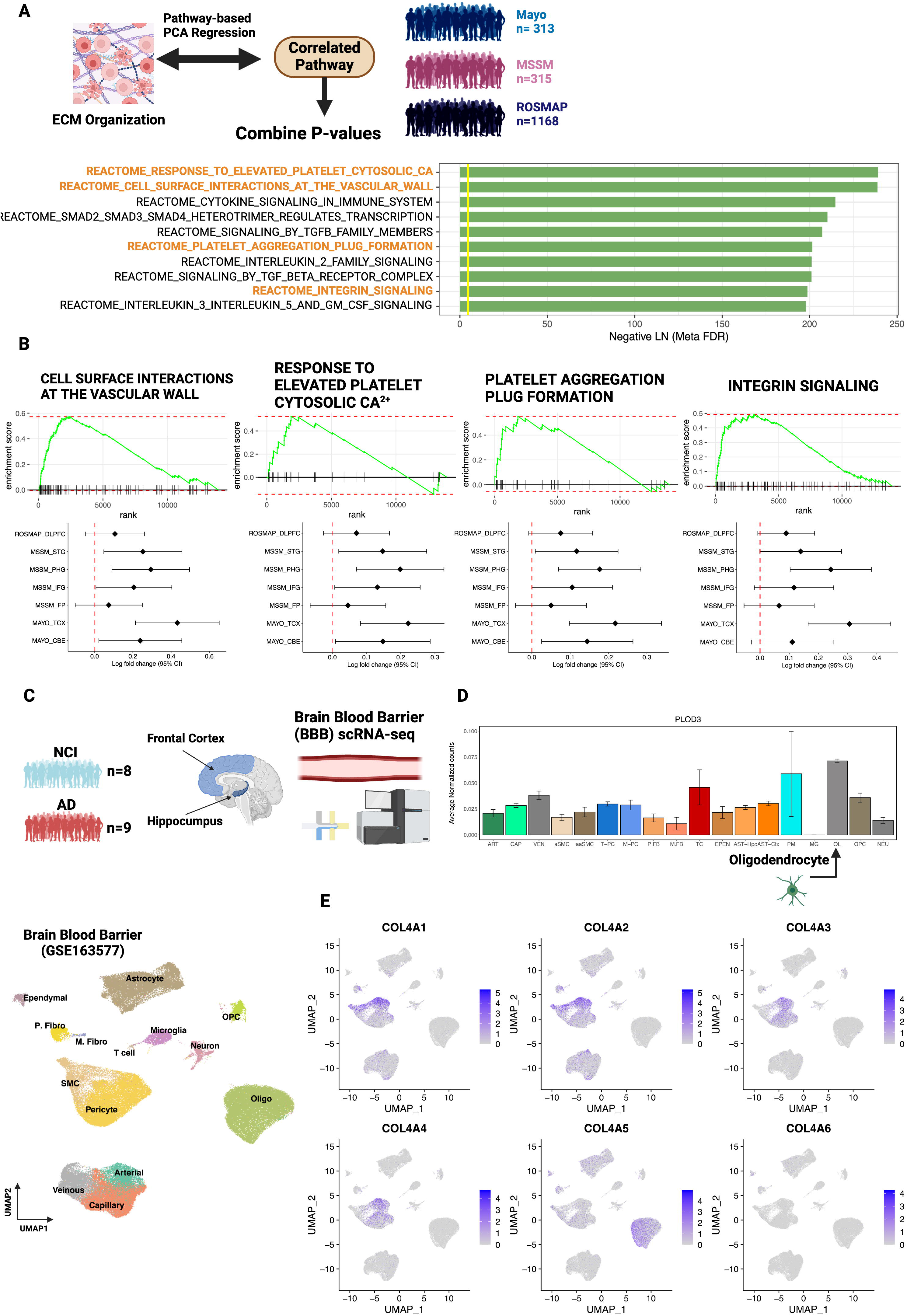
Cerebrovasculature most strongly associated with ECM dysregulation. A Pathways significantly associated with the activity of the ECM organization were estimated for each cohort tissue using the AES-PCA model. The p-values estimated for each cohort and for each brain tissue were estimated as integrated p-values, and the top 10 pathways are shown in the figure. Figures were generated by BioRender.com. B Enrichment of pathways involving the cerebrovasculature in AD with gene set enrichment analysis (GSEA) (FDR < 5%). Forest plots shown below each enrichment plot indicate Log2 fold change for each pathway in each cohort and each region. C Analysis using cerebrovascular scRNA-seq data (8 NCI, 9 AD). D Expression of PLOD3 per cell type. E Expression of type IV collagen per cell type. DLPFC: dorsolateral prefrontal cortex; STG: superior temporal gyrus; PHG: parahippocampal gyrus; IFG: inferior frontal gyrus; FP: frontal pole; TCX: temporal cortex; CBE: cerebellum.

### PLOD3 and COL4A5 are expressed in oligodendrocytes of the cerebrovasculature microenvironment

We analyzed recently reported scRNA-seq data from the vascular microenvironment of the human brain (GSE16357)^45^. These data were used to quantify gene expression by VINE-seq in the cerebral blood vessels in 8 NCI and 9 AD samples (**Figure 5C**). Gene expression was quantified in 143,793 cells from 14 cell types, including vascular endothelial cells (arterial, capillary, and venous), mural smooth muscle cells (SMCs), pericytes, astrocytes, macrophages, T cells, and perivascular and medullary fibroblasts (**Figure 5C**). We examined cell types expressing PLOD3 and COL4A5, which were most strongly expressed in oligodendrocytes (**Figure 5D, E**). In contrast, other type IV collagens were mainly expressed in pericytes and SMCs, which is consistent with the fact that type IV collagen constitutes the vascular basement membrane^46^.

### Oligodendrocytes interact with astrocytes via the COL4A5 ligand

Next, we analyzed the biological functions and pathways mediated by the PLOD3– COL4A5 axis in the cerebrovascular microenvironment. According to the KEGG pathway analysis, COL4A5 may contribute to cell-to-cell communication via ECM ligand receptors (hsa04512). We analyzed how the PLOD3–COL4A5 axis of oligodendrocytes mediates intercommunication between cell types. CellChat^47^ allows for the estimation of cell–cell interactions for each signaling pathway. We estimated cell–cell interactions based on collagen signaling pathways in the AD group. Oligodendrocytes interacted with astrocytes via the COL4A5 ligand and CD44 receptor (**Figure 6A**). This was verified using NicheNet^48^, another intercellular communication estimation algorithm. Among oligodendrocytes, COL4A5 was identified as one of the most promising candidates (**Figure 6B**). In addition to CD44 identified by CellChat, SDC4, DDR2, ITGB8, and ITGAV were predicted to be astrocyte receptors (**Figure 6B**). These receptors were highly expressed in astrocytes (**Figure 6C**).

**Figure 6.**
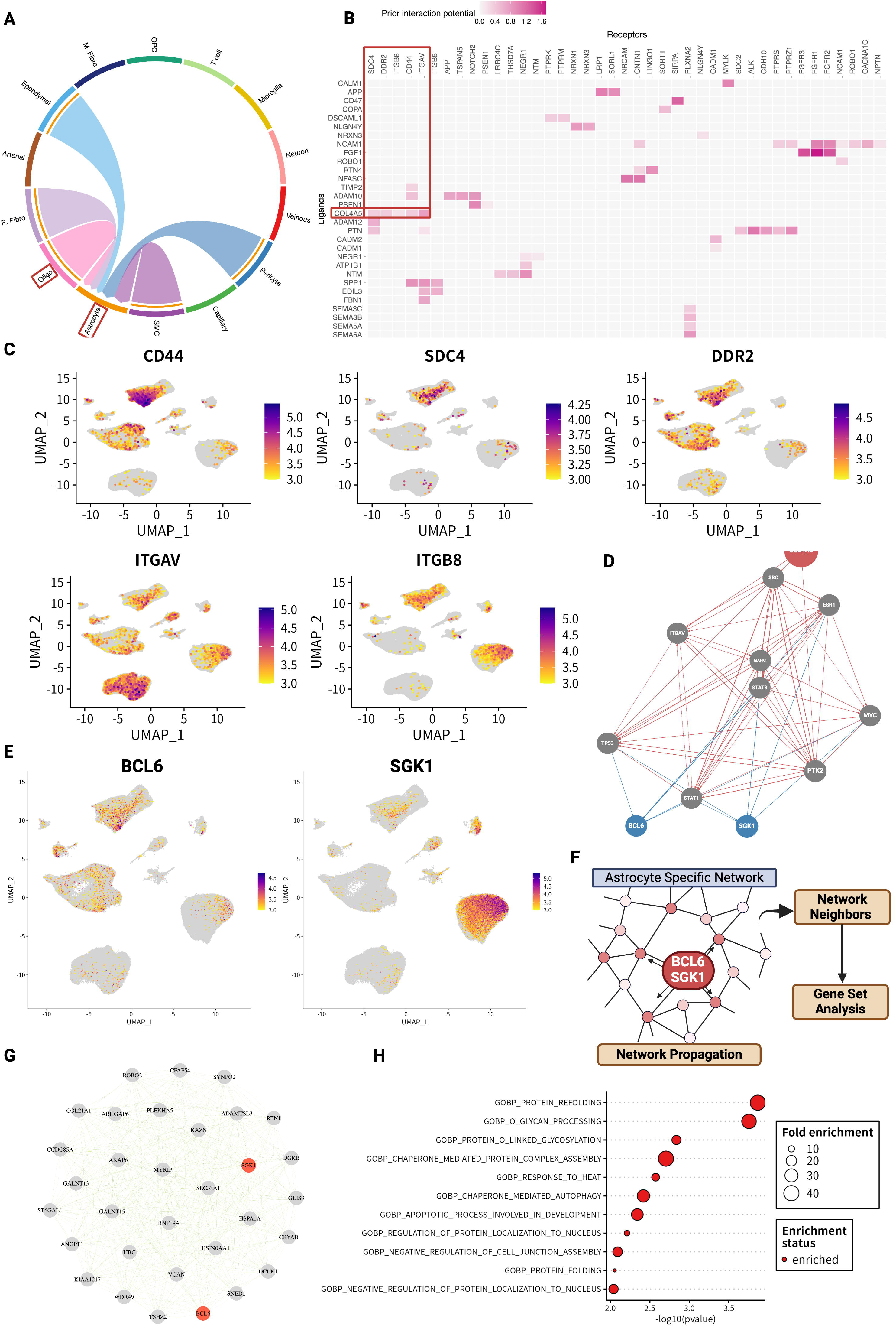
COL4A5 ligand is involved in the regulatory cascade of the astrocyte stress response. A Estimated cell-to-cell communication based on ECM ligand–receptor expression. B Receptor candidates for oligodendrocyte-derived COL4A5 ligands predicted to bind in astrocytes. C Cell type-specific expression levels of receptors for COL4A5. D COL4A5-mediated signaling pathways and target genes in astrocytes. E Expression levels of target genes BCL6 and SGK1 per cell type. F Analysis flow of the exploration of neighboring genes and functional estimation using network propagation in astrocyte-specific co-expression networks. G Top 30 neighboring genes estimated by network propagation based on BCL6 and SGK1. H Gene set analysis of BCL6 and SGK1 neighbor genes.

### COL4A5 ligand is involved in the regulatory cascade of the astrocyte stress response

We performed a detailed analysis of signaling pathways to understand the biological functions of COL4A5-mediated interactions between oligodendrocytes and astrocytes. We integrated the predicted COL4A5 ligand–receptor pairs (CD44, SDC4, DDR2, ITGB8, and ITGAV) into the prior knowledge of the signaling network constructed from multiple perturbation experiments and databases using NicheNet. The results indicated that the COL4A5 ligand targeted and activated B-cell/CLL lymphoma 6 (BCL6) and serum and glucocorticoid-regulated kinase 1 (SGK1) via ECM receptors in astrocytes (**Figure 6D**). BCL6 is a transcription factor and master regulator of humoral immunity and B-cell lymphomagenesis, while SGK 1 encodes a serine/threonine protein kinase that plays an important role in cellular stress responses^49–51^. Both genes were found to be expressed in astrocytes (**Figure 6E**).

Based on these results, we inferred the biological functions of the BCL6 and SGK1 gene modules in astrocytes. An astrocyte-specific co-expression network was constructed based on gene expression using the hdWGCNA algorithm^52^ (**Figure 6F**). Next, we applied the random walk with restart (RWR) algorithm^53^, which is a network propagation algorithm starting from BCL6 and SGK1 on the astrocyte-specific network topology (**Figure 6F**). The RWR allows for the evaluation of the proximity of the network between BCL6, SGK1, and other neighboring genes. Based on these results, we prioritized the top 30 neighbors (**Figure 6G**). GO analysis of these neighboring gene groups revealed that they were enriched mainly for processes involved in stress response (**Figure 6H**). These enriched pathways were also observed in the GO analysis of BCL6 and SGK1 and were independently identified using the network propagation method (**Supplementary Figure 3**).

## Discussion

Our knowledge of the involvement of glycosylation, a major post-translational modification, in the pathogenesis of AD is lacking. We systematically explored the pathogenesis and driving factors based on an integrated analysis of the emerging dimensions of glycosylation in combination with transcriptomics.

In the brain tissue of patients with AD, hyperglycosylation in the ECM is the main signature shared by the glycome and transcriptome, and the glycosyltransferase PLOD3 is an upstream regulator that acts as a functional hub. PLOD3 is predominantly expressed in oligodendrocytes in AD brain tissue and the cerebrovasculature and is co-expressed with COL4A5. Importantly, COL4A5 significantly correlated with APP levels and the activity of the amyloid fiber formation pathway. Single-cell/nuclear analysis revealed that COL4A5 is a ligand for oligodendrocytes that can mediate cell–cell interactions via ECM receptors on astrocytes. In addition, signaling pathway network analysis identified BCL6 and SGK1 as its target genes, and their neighboring genes in the astrocyte-specific network analysis revealed that these two genes are involved in the regulation of the stress response.

The involvement of the ECM in AD has been supported by a large amount of literature^27,54–58^. The physiological roles of the ECM are diverse and include developmental regulation, tissue homeostasis, cell migration, cell proliferation, cell differentiation, neuronal plasticity, and neurite growth^59^. In particular, the ECM is extensively involved in the dysregulation of perineuronal networks in AD^58,60–68^, which are involved in the maintenance of spatial structure, neuronal plasticity, scaffolding,^69^ and the regulation of aggregation; it is also involved in amyloid protein dynamics^27,70–75^ and brain–blood barrier integrity ^41,54,76–79^. As glycoproteins are the major components of the ECM^55,59,80^, glycan synthesis is important for ECM homeostasis in the brain. The enrichment of dysregulated glycoproteins in the ECM is natural in this sense (**Figure 1A, B**).

We discovered that PLOD3 was enriched in the ECM and upregulated in the AD meta-analysis (**Figure 2B–D**). PLOD3 is a multifunctional enzyme, and in addition to its role as a lysyl hydroxylase, it has collagen galactosyltransferase and glucosyltransferase activity^34,81–83^. Although no direct evidence of PLOD3 in AD has been reported, it is known to play an essential role in the formation of collagen, a major component of ECM^84^. For instance, defects in PLOD3 (or lysyl hydroxylase 3; LH3) have been implicated in inherited connective tissue disorders and have been shown to cause cerebral small vessel injury^85,86^, maintenance of the structural integrity of cerebral blood vessels, and the regulation of inflammatory processes^87^. This enzyme is also a promising biomarker of AD, as its expression has been reported to fluctuate in cell-free RNA expression in blood samples from patients with AD^88^.

PLOD3 mediates glycosylation during early collagen formation^31^. Type IV collagen is an essential protein in the cerebral vasculature of patients with AD and is responsible for network formation in the basement membrane. Indeed, in our analysis of single-cell expression levels in cerebral vessels, many type IV collagens (COL4A1/COL4A2/COL4A3/COL4A4) were predominantly expressed in pericytes and SMCs **(Figure 5E)**. In contrast, COL4A5 behaves differently from other type IV collagens and is predominantly expressed in oligodendrocytes. Oligodendrocytes have been shown to stably bind to cerebral blood vessels by zonation analysis based on single-cell/nuclear sequencing analysis^89,90^ and electron microscopy^91^. Interestingly, data from multiple studies support that COL4A5 is strongly correlated with APP and amyloid fiber formation (**Figure 4D**), suggesting a relationship with amyloid plaque accumulation. This may be relevant because the overexpression of type IV collagen generally leads to an increase in cortical basement membrane thickness and has been implicated in the degeneration of cerebral vascular structures^55^. The functional role of type IV collagen in AD cerebrovasculature should be examined in detail in future studies.

We also performed an in silico analysis of cell–cell interactions. COL4A5 functioned as a ligand in oligodendrocyte–astrocyte interactions (**Figure 6A**). Analysis of the signaling pathway network suggested that this cell–cell interaction may contribute primarily to the stress response via SGK1 or BCL6 (**Figure 6D–H**). SGK1 is known to be transcriptionally upregulated under cellular stress^49–51^. On the other hand, both factors have also been reported to be involved in inflammatory responses in the central nervous system. Recent studies have shown that inhibition of SGK1 can suppress the NF-κB-mediated inflammatory pathway in glial cells^92^. There is also evidence that BCL6 plays a central role in regulating astrocytes and NF-κB in response to inflammatory stimuli and disorders^93^. Indeed, in our glycoprotein analysis, the immune response pathway was enriched next to the ECM (**Figure 1B, C**), and inflammatory cytokines were also significantly associated with the ECM organization pathway at the transcriptome level (**Figure 5A, Supplementary Figure S2A**). Inflammatory pathways are key signatures in the AD brain; however, their mechanisms of action in the stress response remain unclear. Further examination of the mechanisms underlying BCL6- and SGK1-mediated stress responses is required.

This study has several limitations. First, the AD glycomic analysis was limited to N-type glycans. Therefore, evidence of ECM hyperglycosylation should be verified in future studies using comprehensive glycoproteomic data. Second, the AD cohort data used in the meta-analysis were limited to those deposited on the AD knowledge portal. To establish a higher level of evidence, data from other large cohort studies should be included. Third, single-cell sequencing data were collected from several different sources; therefore, there is no guarantee that the results reflect the differential expression results of the bulk sequencing used in the meta-analysis. It is expected that this limitation can be overcome in the future as multilayered omics data are collected. However, validation, including experimental approaches, is required.

Our results suggest that glycosylation is involved in the pathogenesis of AD through several unknown mechanisms. Our results also indicate that glycogenomics analysis integrating genetic approaches is a promising method for highlighting the biological functions of glycans and the molecular pathogenesis of diseases at a single-cell resolution. Data on AD glycoproteomics in human subjects are limited. However, as glycoproteomic analysis technology matures, it will be applied to various disease areas, and a vast amount of glycoproteomic data will be accumulated in the next decade. In the near future, the glycogenomics approach will play an important role as a bridge between the established AD genetic pathology and the emerging dimensional omics field of glycoproteomics.

## Methods

### Glycoproteomics enrichment analysis

The first set of glycoproteomics data^9^ was used for enrichment analysis. This dataset was analyzed for glycoproteins overexpressed in the AD group (BRAAK ≥ 5) and the normal group (BRAAK ≤ 2), as defined in the original paper, using the canonical pathway collection of MSigDB (c2.cp.v2022.1.Hs.symbols.gmt). All genes were analyzed as backgrounds using the fedup package in R (https://github.com/rosscm/fedup), and the top 30 significantly enriched pathways were identified. The second set of glycoproteomics data^9^ was analyzed in the same manner. Comparisons were made between the symptomatic group (BRAAK ≥ 5 and CERAD 1 or 2), the asymptomatic group (BRAAK ≥ 3 and CERAD 1 or 2), and the normal group (BRAAK ≤ 2 and CERAD 4), as defined in the original paper. Glycoproteins specifically identified in the symptomatic group were extracted. Enrichment analysis was performed to identify the top 30 significantly enriched pathways.

### Meta-analysis

Meta-analysis using RNA-seq harmonization of AMP-AD followed the published AD-CONTROL analysis protocol (https://github.com/th1vairam/ampad-DiffExp/tree/df3efa793f, 379730bae6d4c9e62910fb2c37e525/gene_level_analysis). First, meta-information was used for data from 3 cohorts (ROSMAP, MSSM, and Mayo), including seven different brain regions, to define patients with definitive late-onset AD from a clinical and neuropathological perspective, that is, neurofibrillary changes, neuritic amyloid plaques, and cognitive dysfunction. The AD control group consisted of patients with AD.

AD controls were defined as patients with few plaques and neurofibrillary changes and no cognitive impairment; in ROSMAP, LOAD cases were those with a BRAAK of 4 or more, a CERAD score of 2 or less, and a cognitive diagnosis of probable AD with no other causes (cogdx = 4); LOAD controls were those with a BRAAK of 3 or less, a CERAD score of 3 or more, and a cognitive diagnosis of “no cognitive impairment” (cogdx = 1). For the MSBB, LOAD cases were defined as those with a CDR score of at least 1, a BRAAK score of at least 4, and a CERAD score of at least 2. LOAD cases were similarly defined as those with a CDR score of 0.5 or less, a BRAAK of 3 or less, and a CERAD of 1 or less as LOAD controls. In Mayo, cases were defined based on neuropathology, with LOAD cases defined as having a BRAAK score of 4 or higher and LOAD controls as having a BRAAK score of 3 or lower.

A meta-analysis using a mixed-effects model was performed to determine the differences in the expression levels of each gene in each of the seven brain regions in each cohort. Effect sizes were estimated using the restricted maximum likelihood method based on the standard mean difference using Hedge. The Metacont function from the meta package of R was used for the analysis. The p-values were corrected for multiple testing by “fdr” using the p.adjust function from the stats package.

### Enrichment map

Gene Set Enrichment Analysis (GSEA) was performed on all genes included in the meta-analysis. The gene set was c2.cp.v2022.1.Hs.symbols from the MsigDB collection, which was loaded using Enrichment Map in Cytoscape and drawn using default settings. After drawing the pathways, we manually classified them into several categories and created several clusters in the network. The list of glycan-related genes manually defined for each glycosylation pathway was then analyzed by post-hoc analysis using the hypergeometric test and the Wilcoxon test, and pathways with FDR ≤ 5% and that were significant by two tests were extracted. Significant pathways in the two tests were extracted.

### Functional hub glycogene identification

Among the enriched pathways based on the same GSEA results as the enrichment map, only pathways containing glycogenes were extracted, and from these, a two-part graph of the pathway–glycogene was extracted. Based on the two-part graphs obtained, each gene was ranked according to its degree of expression. The glycogene with the highest degree was defined as the functional hub glycogene. The results of querying the extracted PLOD3 to the String database (v11) are shown in **Figure 2D**. Forest plots of PLOD3 are shown with estimated effect sizes and 95% confidence intervals from the meta-analysis. For pathway activity, GSEA was performed using the R fgsea package, with gene ranks for effect sizes for each cohort and c2.cp.v2022.1.Hs.symbols for the gene set. Normalized enrichment scores were used for the forest plots.

### Cell-type specificity of PLOD3

For the cell-type specificity of healthy tissues, information was obtained from the Human Proteome Atlas (V22) website by entering the gene name. For data on the entorhinal cortex, information was obtained by entering gene names from http://adsn.ddnetbio.com/.

### Pathway-based PCA regression and GSEA

Pathway-based PCA is a PCA-based method for analyzing pathways and phenotypic associations^43,44,94^. The R Bioconductor PathwayPCA package^42^ was used for the analysis. Using region-specific gene expression data from each AD cohort (Mayo, MSSM, and ROSMAP), we specified the mean expression levels of the ECM pathway component genes as the ECM pathway activity for the objective variable and each pathway other than the ECM pathway for the explanatory variables. The gene set used was c2.cp.v2022.1.Hs.symbols from MSigDB. The pathway names containing “ECM,” “Extracellular,” or “Collagen” were defined as ECM pathways. The genes involved in these pathways were defined as signatures.

The p-values of the list of pathways significantly associated with ECM were combined using Fisher’s method to calculate the integrated p-value. For the calculation, the log-sum function of the R metapackage^95^ was used, and the p-values of the individual datasets were entered for each pathway. In addition, we cross-checked whether significantly related pathways were sufficiently enriched at the expression level. Focusing on the top 10 pathways, we applied GSEA based on the gene set c2.cp.v2022.1.Hs.symbols from MSigDB using the effect sizes of the 3 cohort meta-analysis as the gene rank. To further validate that the top 10 pathway activities tended to increase by cohort and region, the means of the effect sizes and confidence intervals were calculated for the signature genes and illustrated as forest plots.

### Analysis of brain vasculature with scRNA-seq

Count data were preprocessed using the Seurat package in R. Normalization, feature selection with VST, scaling, and dimensional reduction using PCA and UMAP were performed. The cell types were visualized using those previously identified in an original paper^45^. Next, for each cell type, variation analysis between the AD and cognitively normal groups was performed using Seurat’s FindMarkes function, and enrichment analysis for the identified groups of DEGs was performed using the R fedup package. The c2.cp.v2022.1.Hs.symbols from MSigDB was used as the gene set to determine which cell types were enriched in ECM-related pathways. We selected gene sets with pathway names containing “ECM,” “Extracellular,” “Matrisome,” or “Collagen” in the pathway name. The enriched p-values were further transformed as -Log10(FDR) from the multiple-test-corrected FDR, considered as differentially expressed activity signals, and visualized using a heatmap. The expression levels per cell type were obtained by querying https://twc-stanford.shinyapps.io/human_bbb/ for PLOD3.

### Cell–cell interaction and signaling network analysis

Cell–cell interactions were analyzed using the R package CellChat^47^ (https://github.com/sqjin/CellChat). Oligodendrocytes and astrocytes identified with CellChat were further analyzed using another algorithm, NicheNet^48^ (https://github.com/saeyslab/nichenetr). For a detailed analysis, the ligand–receptor prior information was input by integrating the ligand–receptor pair information used in CellChat with the ligand–receptor pair information used in NicheNet. This was also used for the signal network analyses. A pre-built model was downloaded (https://github.com/saeyslab/nichenetr/blob/master/vignettes/model_construction.md), and both the ligand–receptor information and expression information were identified in the cell– cell interactions.

### Astrocyte cell-type specific network propagation

For astrocyte-specific network construction using cerebrovascular scRNA-seq, the Toplogical Overlap Measure (TOM) was estimated using hdWGCNA^52^, and edges were further defined only if they had a TOM above the 90th percentile as a threshold. The network propagation method was applied using the R package RandomWalkRestartMH^53^. In other words, we performed an RWR starting from SGK1 and BCL6 in the obtained network topology. The 30 most relevant neighbors were narrowed down and plotted using the R package igraph. The R package fedup was used for the enrichment analysis. To estimate the transcriptional activity of BCL6, curated regulon information was first obtained using the R package DoRothEA^96^, and transcription factor target genes were estimated using the Viper^97^ algorithm. The R package decoupleR^98^ was used for the analysis.

## Acknowledgements

This work was supported by the Human Glycome Atlas Project (HGA) and JSPS KAKENHI, Grant Number: JP20H04282. The results published herein are partly based on data obtained from the AD Knowledge Portal (https://adknowledgeportal.org). Data generation was supported by the following NIH grants: P30AG10161, P30AG72975, R01AG15819, R01AG17917, R01AG036836, U01AG46152, U01AG61356, U01AG046139, P50 AG016574, R01 AG032990, U01AG046139, R01AG018023, U01AG006576, U01AG006786, R01AG025711, R01AG017216, R01AG003949, R01NS080820, U24NS072026, P30AG19610, U01AG046170,

RF1AG057440, and U24AG061340, as well as the Cure PSP, Mayo, and Michael J Fox foundations, Arizona Department of Health Services, and the Arizona Biomedical Research Commission. We thank the participants of the Religious Order Study and Memory and Aging projects for their generous donations, the Sun Health Research Institute Brain and Body Donation Program, Mayo Clinic Brain Bank, and Mount Sinai/JJ Peters VA Medical Center NIH Brain and Tissue Repository. Data and analysis contributing investigators included Nilüfer Ertekin-Taner, Steven Younkin (Mayo Clinic, Jacksonville, FL), Todd Golde (University of Florida), Nathan Price (Institute for Systems Biology), David Bennett, Christopher Gaiteri (Rush University), Philip De Jager (Columbia University), Bin Zhang, Eric Schadt, Michelle Ehrlich, Vahram Haroutunian, Sam Gandy (Icahn School of Medicine at Mount Sinai), Koichi Iijima (National Center for Geriatrics and Gerontology, Japan), Scott Noggle (New York Stem Cell Foundation), and Lara Mangravite (Sage Bionetworks).

## Author contributions

**Yusuke Matsui:** Conceptualization, Methodology, Software, Validation, Formal analysis, Investigation, Resources, Data Curation, Writing - Original draft, Writing - Reviewing and Editing, Visualization, Funding acquisition. **Akira Togayachi**: Resources, Data Curation, Writing - Reviewing and Editing **Kazuma Sakamoto**: Writing - Reviewing and Editing **Kiyohiko Angata**: Writing - Reviewing and Editing. **Kenji Kadomatsu**: Supervision, Writing-Reviewing and Editing, Project administration, Funding acquisition. **Shoko Nishihara:** Conceptualization, Supervision, Writing-Reviewing and Editing, Project administration.

## Data availability

The glycoproteomics datasets were compiled and used from the supplementary files published in the respective papers^9,10^. For the AMP-AD transcriptome dataset, data were obtained from the RNAseq Harmonization Study (RNAseq Harmonization) Repository (syn21241740). Single-cell transcriptome data in the entorhinal cortex were obtained from GSE138852. Brain vascular data were downloaded from GSE163577.

---

**Supplementary Figure 1.**
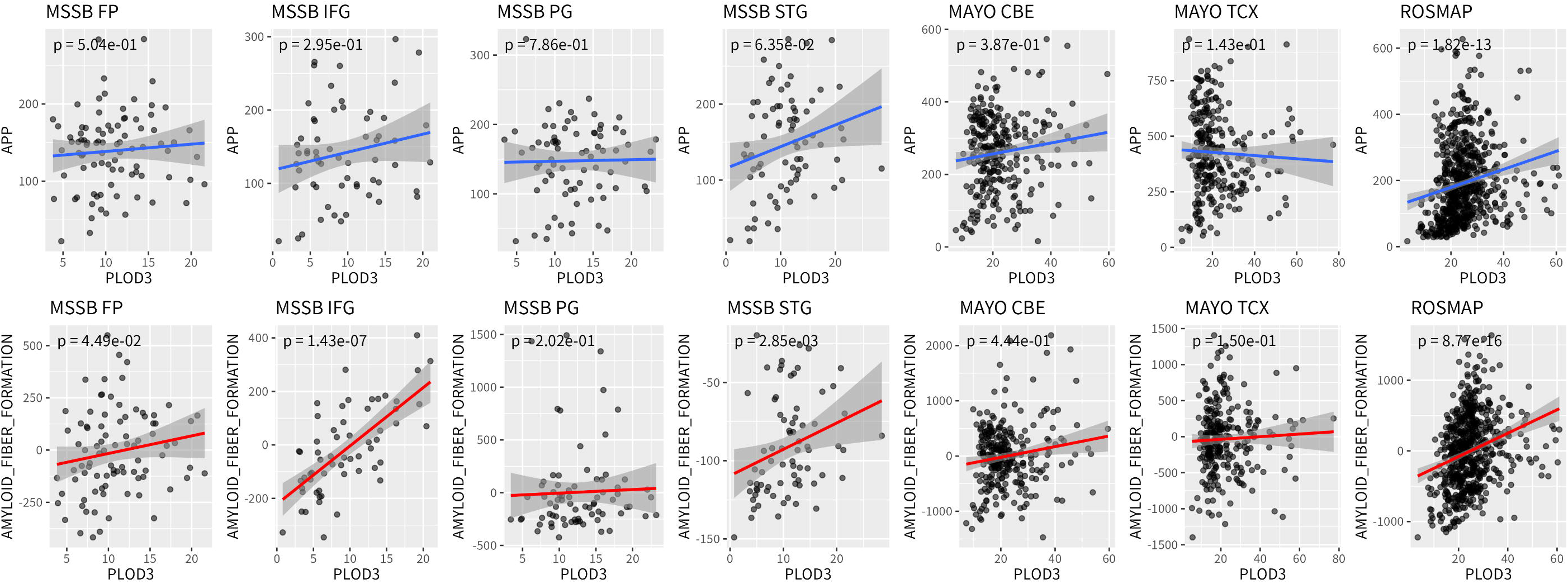
Correlation of PLOD3 with APP activity (upper panel) and amyloid fiber formation (lower panel) for each cohort and each region. DLPFC: dorsolateral prefrontal cortex; STG: superior temporal gyrus; PHG: parahippocampal gyrus; IFG: inferior frontal gyrus; FP: frontal pole; TCX: temporal cortex; CBE: cerebellum.

**Supplementary Figure 2.**
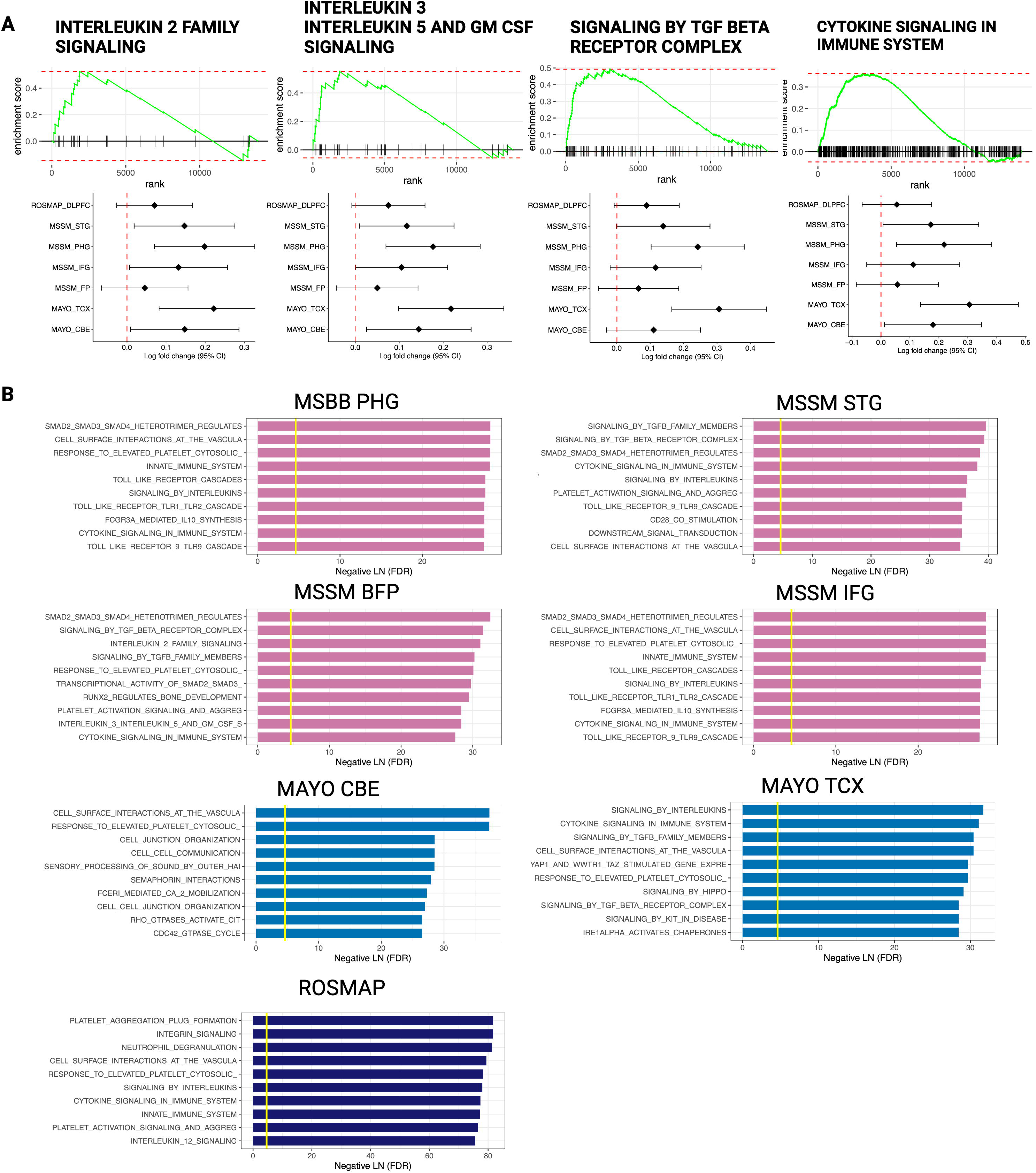
ECM activity is strongly associated with inflammatory cytokines. A Enrichment of pathways involving the immune system in AD with GSEA (FDR < 5%). Forest plots shown below each enrichment plot indicate Log2 fold change for each pathway in each cohort and each tissue. B Pathways significantly associated with ECM activity obtained by applying AES-PCA for each cohort and region. DLPFC: dorsolateral prefrontal cortex; STG: superior temporal gyrus; PHG: parahippocampal gyrus; IFG: inferior frontal gyrus; FP: frontal pole; TCX: temporal cortex; CBE: cerebellum.

**Supplementary Figure 3.**
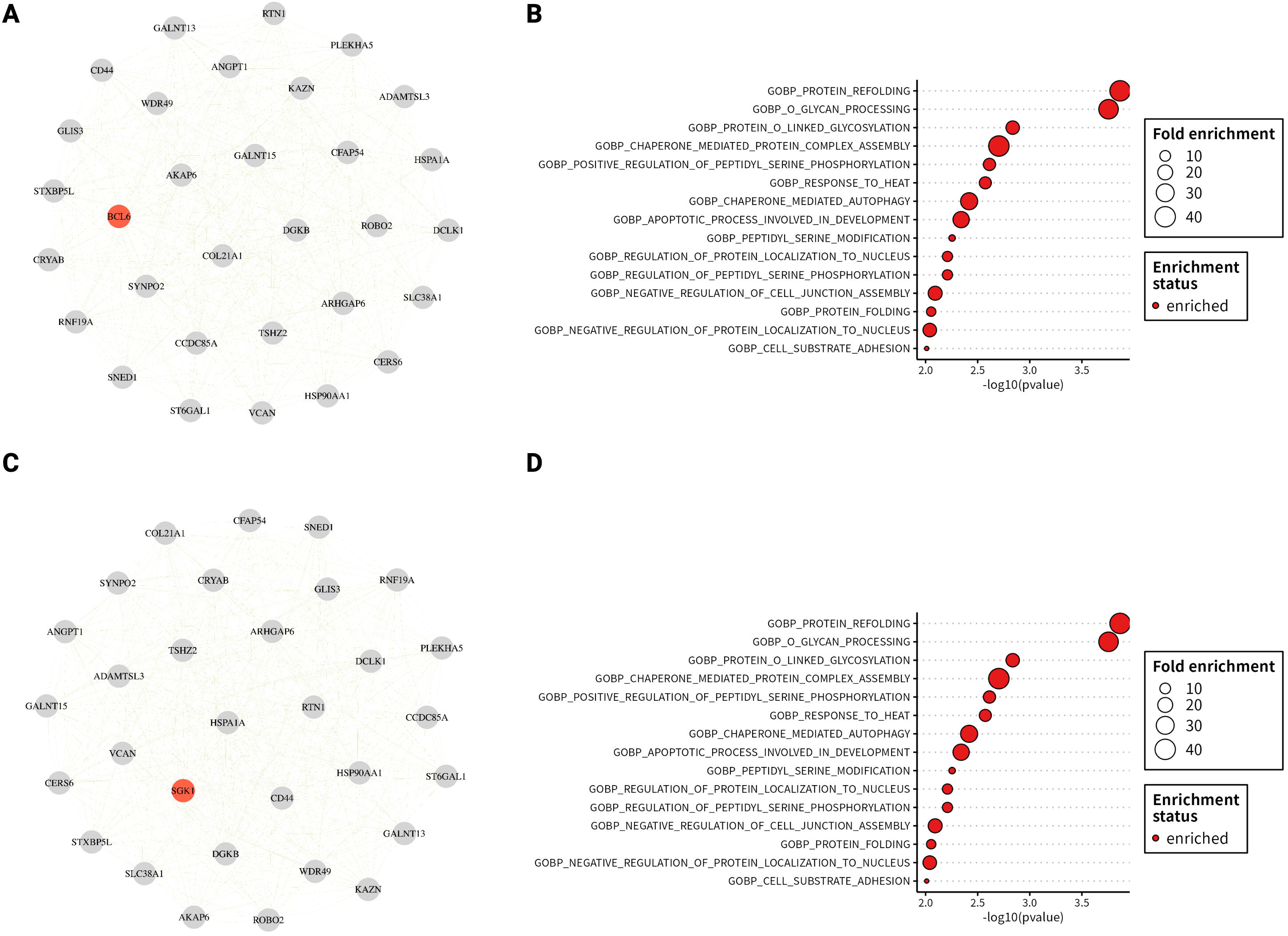
COL4A5 ligand is involved in the regulatory cascade of the astrocyte stress response. A Top 30 neighboring genes estimated by network propagation based on BCL6. B Gene set analysis of BCL6 neighbor genes. C Top 30 neighboring genes estimated by network propagation based on SGK1. D Gene set analysis of SGK1 neighbor genes.

